# Intermittent fasting has a diet-specific impact on the gut microbiota and colonic mucin *O*-glycosylation of mice

**DOI:** 10.1101/2022.09.15.508181

**Authors:** Hasinika K.A.H. Gamage, Abdulrahman M. M. Sathili, Krishnatej Nishtala, Raymond W.W. Chong, Nicolle H. Packer, Ian T. Paulsen

## Abstract

The colonic mucus layer and microbiota adhered to it are vital for mediating host metabolic, immune, and gut health. Yet, how intermittent fasting impacts these microbial communities and *O*-glycosylation of mucin proteins, the predominant component of the colonic mucus layer, remains largely unexplored. Here, using a C57BL/6J mouse model fed either a high-fat diet or normal chow, we examined the impact of a two-day a week fasting regimen on host physiology, faecal and colonic mucosal microbiota, and mucin *O*-glycosylation. Our results demonstrated distinct diet-specific impacts of intermittent fasting on host physiology; mice fed the high-fat diet had a lower body weight and improved glucose tolerance upon fasting, whilst there were no significant changes in mice fed the normal chow. This was observed despite the similar feed and energy intake between groups with and without fasting. There were significant changes in the faecal and colonic mucosal microbiota community structure and composition, and mucin *O-*glycosylation upon fasting in both dietary groups, but the specific nature of these alterations was diet-dependent. Correlation analysis revealed significant associations between fasting-mediated changes in the abundance of specific mucosal bacteria and *O*-glycan structures. While intermittent fasting is a popular means of extending healthy life expectancy, there is a lack of information on its impacts on the mucosal microbiota and colonic mucus layer, which are key determinants of gut health. Our study addresses this knowledge gap and serves as the first report on how intermittent fasting influences colonic mucin *O*-glycosylation and the associations between mucosal glycans and bacteria.

## Introduction

Frequent and abundant consumption of food, particularly those high in fat and sugar is associated with perturbations to the gut microbiota and the onset and progression of conditions such as obesity and metabolic disease [1-3]. Intermittent fasting through periodic dietary restriction is an increasingly popular means for ameliorating these changes. Fasting without a reduction in overall caloric intake has been shown to promote optimal energy metabolism, alleviate symptoms of metabolic syndrome and extend the healthy lifespan of various animal models [4-8]. Specific intermittent fasting regimens have also been studied for their ability to improve gut health [8, 9] and influence gut microbial populations and colonic mucus layer.

The gut microbiota plays an essential role in maintaining gut health, host metabolism and immunity; changes to the composition and functions of gut microbial communities can have profound impacts on host health. Previous studies have shown a clear impact of fasting on the gut microbiota [9-11]. For example, time-restricted feeding in mice has been linked with an increase in the abundance of *Oscillibacter* and *Ruminococcaceae*, and a decrease in *Lactobacillus* in the gut microbiota compared to that of mice fed *ad libitum* [9]. Intermittent fasting has also been shown to ameliorate high-fat diet induced changes in the composition and diurnal oscillations in the gut microbiota of mice [9, 12]. In fact, the gut microbiota has been proposed to play a key role in facilitating the beneficial effects of fasting on host health [6, 13, 14]. A resistance to intermittent fasting-induced browning of white adipose tissue, which is linked to impaired energy expenditure and obesity, has been reported in mice with a depleted gut microbiota [6]. Transplanting these mice with the gut microbiota of healthy mice subjected to the same fasting regimen activated white adipose browning, indicating a gut microbiota mediated impact of intermittent fasting on fat tissue browning and metabolic activities [6].

The colonic mucus layer, which is located at the interface between the gut microbiota and colonic lumen, plays an integral role in maintaining microbiota-host homeostasis and gut health [15, 16]. Previous examinations of how intermittent fasting influences the colonic mucosa have demonstrated significant improvements in the integrity and thickness of the mucus layer, particularly in conditions such as ulcerative colitis and enteric infections [17-19]. Fasting has also been associated with an increase in the total gastric mucosal glycoproteins [20], the main component of the mucus layer. Mucosal glycoproteins, known as mucins, belong to a family of highly glycosylated proteins heavily decorated with *O*-linked glycans [21]. Mucin *O*-glycans in the colon serve as cell adhesion sites and provide an alternative nutritional source for the mucosal microbiota [22]. The interactions between colonic mucin *O*-linked glycans and microbiota are key to the development of the mucus layer and microbial colonisation in the gut [21, 23], and are vital for regulating gut and host health [15]. While changes in mucin *O*-glycosylation could have profound impacts on the gut microbiota and host [15, 16], the impact of intermittent fasting on colonic mucin *O*-linked glycans remains unexplored.

Dietary composition, in addition to meal frequency is a key modulator of both the gut microbiota and colonic mucin *O*-glycosylation. Previous studies have demonstrated clear changes in the abundance of specific gut bacteria and mucin *O*-glycans upon dietary changes, particularly with changes in dietary fibre intake [24, 25]. Furthermore, diet has been associated with changes in how the gut microbiota interact with mucin *O*-glycans [24]. While examining the effects of both fasting and composition of diet on non-fasting days is crucial for understanding the impact of intermittent fasting regimens on gut microbial communities and mucin *O*-glycosylation, there is a lack of direct comparative studies on this. A study limited to examining only the host health has shown distinct impacts of a time-restricted feeding regimen on the expression levels of hepatic circadian oscillator components in mice fed a normal chow and high-fat diet [5]. However, there is limited information on this in relation to the gut microbiota and colonic mucin *O*-glycosylation.

In this study, we examined the impact of an intermittent fasting regimen on C57BL/6J mice fed either a high-fat diet or normal chow, two diets with distinct compositions. The impact of the dietary regimens on the colonic mucosal and faecal microbiota, colonic mucin *O-* glycosylation, and host metabolic health was investigated.

## Results and Discussion

### Intermittent fasting had a diet-specific impact on host physiology

C57BL/6J mice were fed either a high-fat diet or a normal chow for 12 weeks with *ad libitum* access or were subjected to an intermittent fasting regimen with no access to feed on two non-consecutive days a week followed by *ad libitum* feeding on the remaining days (Figure S1). Mice fed the high-fat diet *ad libitum* (HF) had a significantly higher body weight, (Fig. 1A and Figure S2A), overall body weight gain (Figure S2B) and epididymal fat pad mass (Figure S3A) compared to that in mice fed the normal chow *ad libitum* (NC). This potentially suggests the development of diet-induced obesity in the HF group. All these measurements were significantly lower in the high-fat diet with intermittent fasting (HF-IF) group compared to the HF group; this was observed despite the similar amounts of food (Fig. 1B) and energy (Figure S2C) intake between the two groups, likely indicative of the lower metabolic efficiency and/or higher energy expenditure in mice upon fasting. Compared to the NC group, mice fed the normal chow with intermittent fasting (NC-IF) showed no significant reduction in body weight or epididymal fat mass, suggesting a lower impact of the fasting regimen on mice fed the normal chow. Both the HF-IF and NC-IF groups demonstrated a reduction in the body weight immediately after each fasting cycle, but this was recovered within less than 24 hours when *ad libitum* access to feed was restored (Figure S2A).

**Figure 1.**
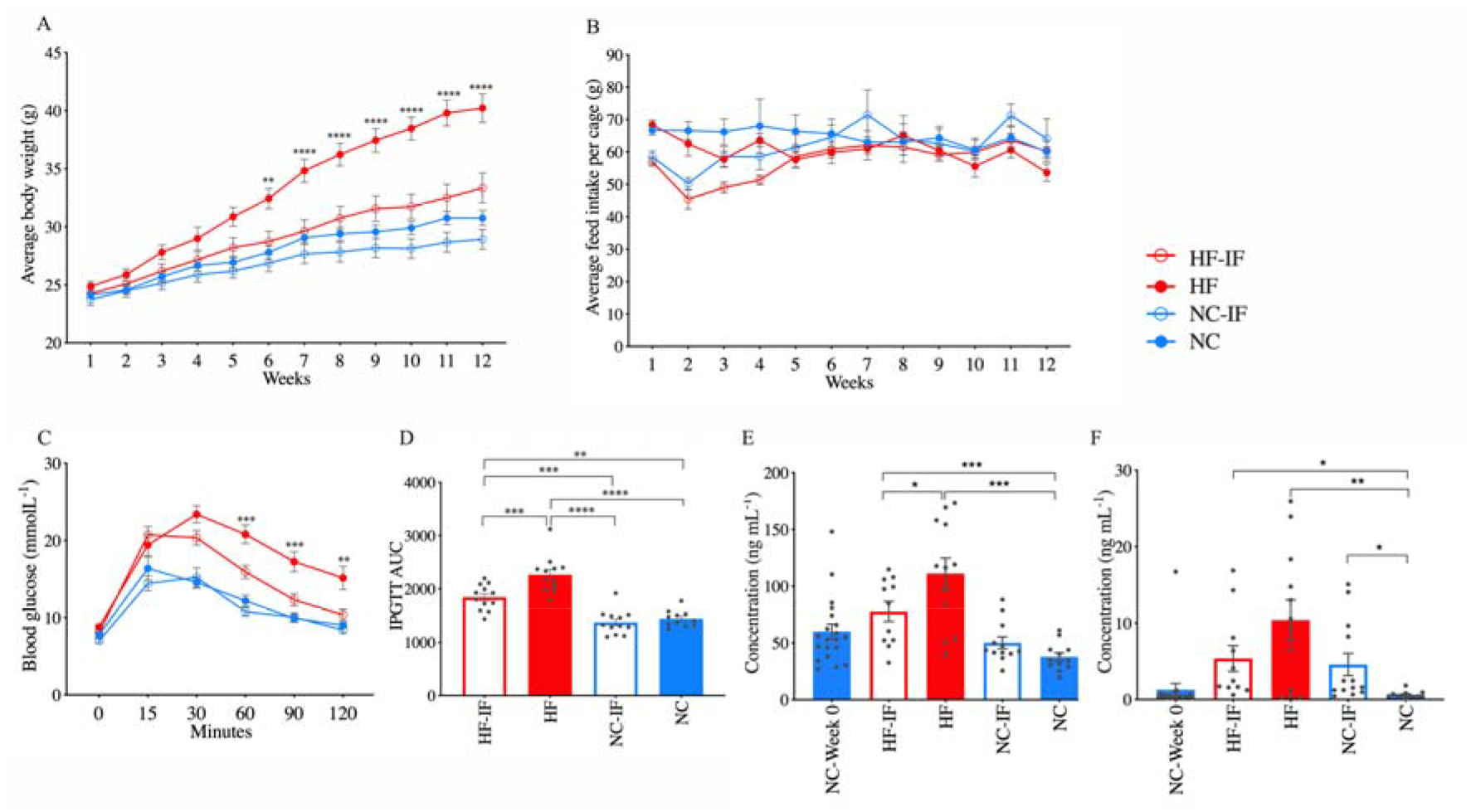
Impact of intermittent fasting on host physiology. Weekly measurements of **(A)** body weight per mouse and **(B)** feed intake per cage are shown with the **(C)** blood glucose levels and **(D)** area under the curve (AUC) of the intraperitoneal glucose tolerance test (IPGTT) conducted at week 4 and blood plasma concentrations of **(E)** resistin and **(F)** leptin at week 0 and 12. Data are shown as mean ± SD for each treatment group: high-fat diet with intermittent fasting (HF-IF) or *ad libitum* (HF) and normal chow with intermittent fasting (NC-IF) or *ad libitum* (NC). Significance was determined based on two-way ANOVA with Tukey’s multiple comparison tests or Kruskal-Wallis test with Dunn’s multiple comparison tests where appropriate (**** *P* < 0.0001, *** *P* < 0.001, ** *P* < 0.01 and * *P* < 0.05).

We then examined the impact of fasting on specific indicators of host metabolic health. Intraperitoneal glucose tolerance tests (IPGTTs) conducted at week 4 and 11 revealed a significantly higher ability in the HF-IF group to regulate glucose levels compared to that in the HF group, with no difference observed between the NC and NC-IF groups (Fig. 1C and 1D, Figure S2D and S2E). Despite having similar liver masses (Figure S3C), the HF-IF group demonstrated lower levels of liver steatosis compared to the HF group (Figure S3D), suggesting an intermittent fasting-induced reduction in the build-up of fat in the liver. Upon examination of specific cytokines and biomarkers of obesity and diabetes (Fig. 1E, 1F and Figure S4), we observed the highest concentration of resistin and leptin in the blood plasma of mice in the HF group (Fig. 1E and 1F). Whereas a significant reduction of both biomarker levels was seen in the HF-IF group. In the NC-IF group, leptin levels were significantly higher than that in the NC group, with no significant difference in resistin levels. Resistin expression has been previously shown to be higher with elevated blood glucose levels and obesity [26, 27], which is consistent with our observations. A diet-specific change in leptin levels was observed upon fasting; there was an increase in leptin concentrations in the NC-IF group, while a drop was observed in the HF-IF group. Leptin regulates host energy balance and contributes to metabolic adaptation in response to caloric restriction [28]. Lower leptin levels have been linked with an increase in appetite and *vice versa* [29]. While we observed differences in plasma leptin concentrations, there were no significant differences in weekly feed intake between groups (Fig. 1B). Circulating levels of leptin and sensitivity to it have been shown to depend on multiple factors including body weight [29], a likely explanation for the observed diet-specific differences in plasma leptin levels upon fasting.

Intermittent fasting regimens typically consist of periods with no or low caloric intake followed by *ad libitum* feeding on the remaining days [2]. While diet on non-fasting days could be critical for the efficacy of fasting, there is a lack of direct examinations of this. Our results suggest a significant impact of dietary composition on the outcomes of intermittent fasting. The two tested diets, high-fat diet and normal chow have distinct nutritional compositions (Table S1), including significantly different total fat contents, 23.5% and 4.8%, respectively. A previous study with time-restricted feeding has also reported greater improvements in the body weight and host circadian oscillation in C57BL/6J mice fed a high-fat diet compared to a normal chow [5]. These observations suggest intermittent fasting is particularly effective in ameliorating high-fat diet-induced changes in host metabolic health. Although this 12-week intermittent fasting regimen tested in our study has a minimal impact on the metabolic health of mice fed a normal chow, it could have longer-term effects, such as a longer lifespan as previously reported with caloric restriction [30]. Future life-long experiments will be essential in elucidating whether the tested feeding regimen is linked with a higher life expectancy.

### Fasting induced diet-specific changes in the mucosal and faecal microbiota

The impact of fasting on the mucosal and faecal microbiota was determined through sequencing of the 16S rRNA gene amplicons. The overall microbial communities in the mucus and faecal samples showed significant shifts in mice fed the high-fat diet compared to that of mice fed the normal chow (*P* = 0.0001, PERMANOVA, Fig. 2A and 2D), showing a dramatic effect of dietary composition on the microbiota community structure. Intermittent fasting significantly shifted (*P* < 0.05) the overall structure of the mucosal and faecal microbiota of mice in the HF-IF and NC-IF groups. In the HF-IF group, the shift in the faecal microbiota (*P* = 0.0008) was more prominent compared to that in the mucosal microbiota (*P* = 0.02). Conversely, the NC-IF group demonstrated greater changes in the mucosal microbiota community structure (*P* = 0.0001) than the faecal microbiota (*P* = 0.05). This observation suggests a diet- and body site-specific impact of fasting on the community structure of faecal and mucosal microbiota. The alpha diversity, as determined by Shannon diversity and Simpson’s evenness indices, was significantly higher in the NC group compared to the HF group, with no significant change upon fasting (Fig. 2). Similar to our observations, a lack of changes in the microbiota alpha diversity upon specific fasting or caloric restriction regimens has been previously reported [8, 9], which highlights a stronger impact of dietary composition in shaping the overall gut microbiota diversity compared to meal frequency.

**Figure 2.**
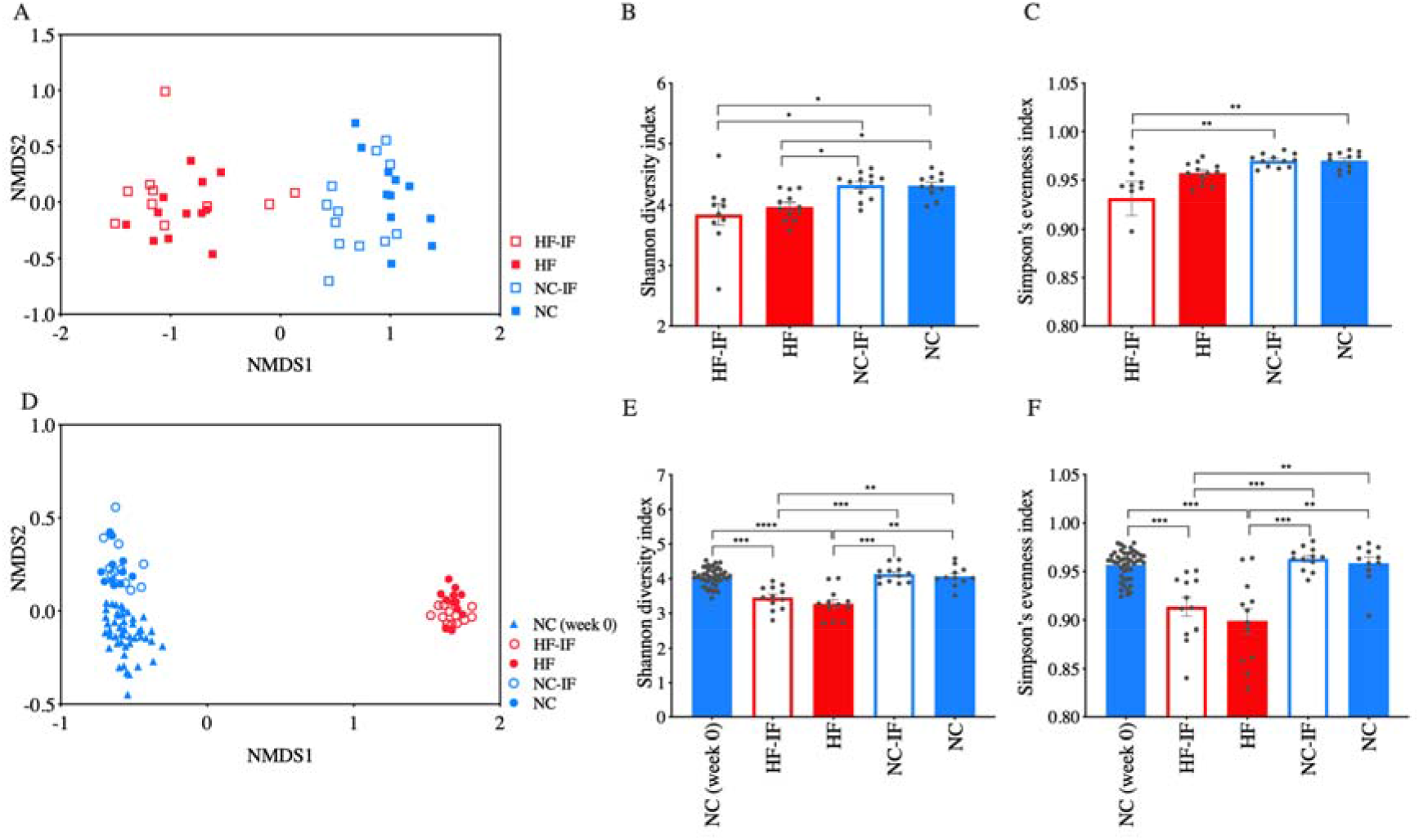
Ordination and alpha diversity of the mucosal and faecal microbiota. **(A)** Ordination of the mucosal microbiota at week 12. Mucosal microbiota alpha diversity shown as **(B)** Shannon diversity and **(C)** Simpson’s evenness indices. **(D)** Ordination of the faecal microbiota at weeks 0 and 12. **(E)** Shannon diversity and **(F)** Simpson’s evenness indices of the faecal microbiota samples collected at week 0 and 12. Microbiota community structure of samples is shown as Bray-Curtis similarity based non-metric multidimensional scaling (nMDS) plots. Alpha diversity is shown as mean ± SD for each treatment group: high-fat diet with intermittent fasting (HF-IF) or *ad libitum* (HF) and normal chow with intermittent fasting (NC-IF) or *ad libitum* (NC). Significance was determined using Kruskal-Wallis test with Dunn’s multiple comparison tests (**** *P* < 0.0001, *** *P* < 0.001, ** *P* < 0.01 and * *P* < 0.05).

To examine the impact of fasting on the abundance of microbial species equivalents, demonstrated as amplicon sequence variants (ASVs), we conducted linear discriminant analysis effect size (LEfSe) comparisons between treatment groups. These identified significant changes in the abundance of specific microbes in the mucosal (Fig. 3) and faecal (Figure S5) microbiota of mice in the HF-IF and NC-IF groups compared to the HF and NC groups, respectively. The abundance of some bacteria changed similarly across dietary groups and body sites upon fasting; ASVs in the genus *Lactobacillus* and family *Ruminococcaceae* UCG-013 were highly abundant in both the mucosal and faecal microbiota of the HF-IF (Fig. 3A and Figure S5A) and NC-IF (Fig. 3B and Figure S5B) groups, demonstrating clear impacts of fasting irrespective of the composition of diets. Previous studies have also reported an increase in the abundance of *Lactobacillus* and *Ruminococcaceae* with caloric restriction [8, 31] and time-restricted feeding [9], respectively. A higher abundance of these bacteria has been linked with improved metabolic health [32], and *Lactobacillus* has been associated with a longer lifespan in mice [33]. Our data demonstrate a higher *Lactobacillus* and *Ruminococcaceae* abundance in both the HF-IF and NC-IF groups with significant improvements in the tested metabolic health parameters observed only in the HF-IF group. This may indicate a diet-specific association of these two bacterial groups with the metabolic health of the host. Future life-long intermittent fasting experiments will be useful in gaining further insight into this, especially, in understanding the role of these bacteria when a low-fat and high-fibre diet is consumed.

**Figure 3.**
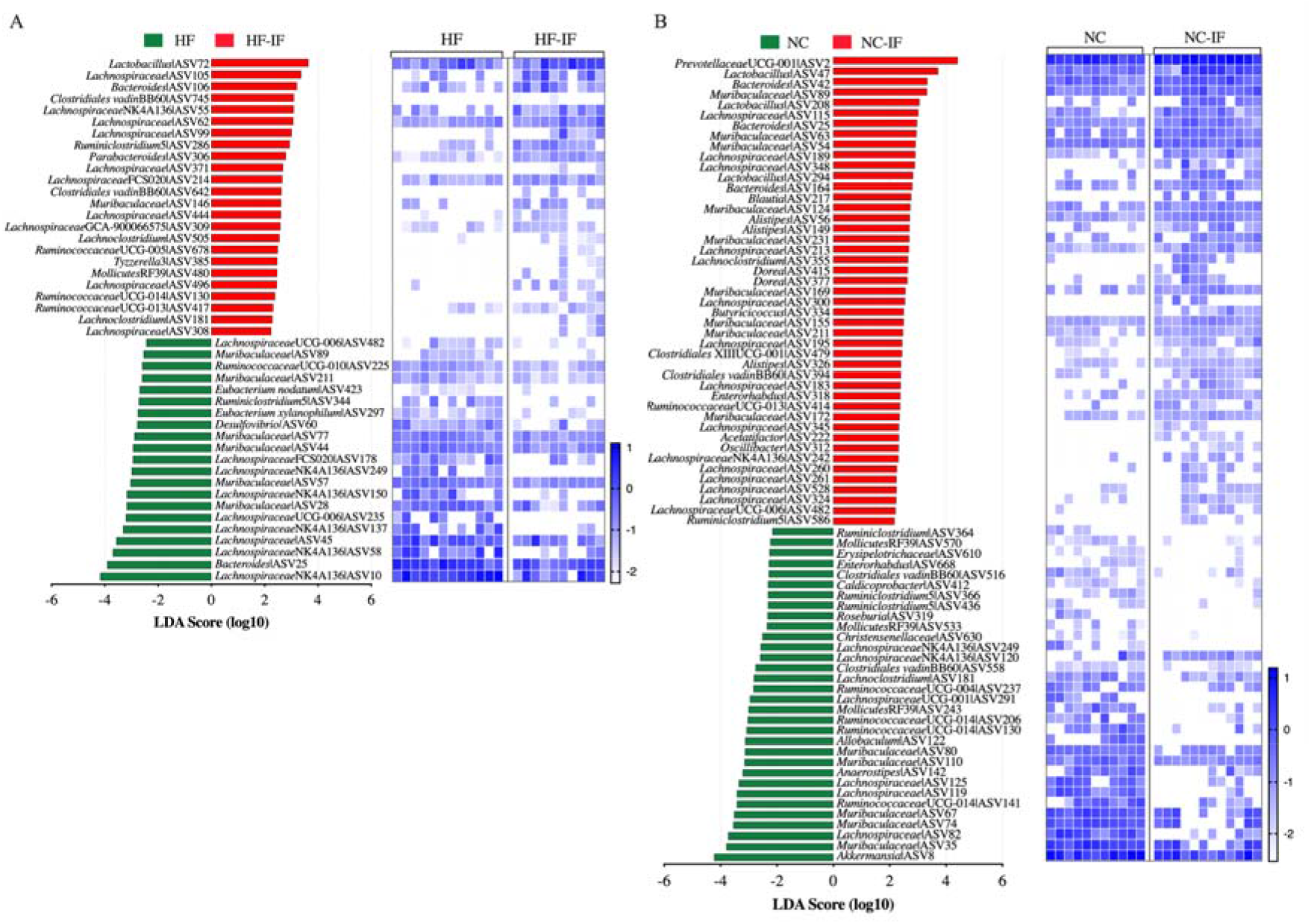
Effect of intermittent fasting on amplicon sequence variants (ASVs) in the colonic mucosal microbiota at week 12. Data were obtained using the linear discriminant analysis effect size (LEfSe) method through comparing the abundance of ASVs between treatment groups: **(A)** high-fat diet *ad libitum* vs intermittent fasting (HF vs HF-IF) and **(B)** normal chow *ad libitum* vs intermittent fasting (NC vs NC-IF). The histograms show linear discriminant analysis (LDA) scores computed for each ASV. The unpaired heatmaps show the relative abundance (Log_10_ transformed) of the ASVs, rows correspond to the abundance of the ASVs, and columns correspond to individual mice in each dietary group. Blue and white denote the highest and lowest relative abundance, respectively, as shown in the legend.

Most fasting-associated changes in the mucosal and faecal microbial communities were diet specific. For example, the mucosal microbiota of mice fed the HF-IF had a higher abundance of ASVs in *Lachnoclostridium* compared to the HF group. Whereas, in the NC-IF group, the abundance of ASVs in genera such as *Bacteroides* and *Alistipes* was higher, while the abundance of *Akkermansia* was lower compared to the NC group (Fig. 3). Concurrently, in the faecal microbiota of the HF-IF group, the abundance of ASVs in *Lachnospiraceae* was higher, while the abundance of *Rikenellaceae* and *Christensenellaceae* was lower (Figure S5). Changes in the abundance of these bacterial groups have been linked with host metabolic and immune health [34, 35]. For example, *Alistipes* has been shown to provide protective effects against diseases, including liver fibrosis and colitis [34], while a reduction of *Christensenellaceae* has been reported with weight loss [36].

Taken together, our findings suggest a substantial impact of intermittent fasting on the gut microbiota. While most microbiota changes were specific to either the HF-IF or NC-IF group, some changes were observed in both groups, demonstrating both diet-dependent and independent impacts of fasting on the gut microbiota composition. Some high-fat diet-induced shifts in the microbiota were also altered with fasting. Consumption of the high-fat diet *ad libitum* resulted in vastly different mucosal and faecal microbiota community structures (Fig. 2) and compositions (Table S2) compared to the NC group, this was consistent with previous reports on alterations in the gut microbiota upon consumption of a high-fat diet [24, 37, 38]. While fasting did not reverse all high-fat diet-induced modifications, it changed the abundance of specific gut bacteria, including ASVs in the families *Christensenellaceae, Lachnospiraceae* and *Ruminococcaceae*, which have been previously linked with the intake of a high-fat diet and obesity in mice [24, 38]. To our knowledge, this is the first investigation on how intermittent fasting effects the mucosal microbiota structure and composition and the impact of dietary composition in mediating these changes. Observed diet-specific changes in the faecal and mucosal microbiota warrants more detailed future studies into understanding the role of dietary composition on non-fasting days on the efficacy of intermittent fasting regimens.

We then compared the colonic mucosal and faecal microbiota and identified significant differences not only in the microbial composition (Figure S6) but in the precise nature fasting influenced the microbiota of the two body sites (Fig. 3 and Figure S5). For example, the mucosal microbiota had a higher abundance of ASVs belonging to *Lachnospiraceae, Muribaculaceae* and *Lactobacillus* across all four treatment groups compared to that in the faecal samples, consistent with previous reports [39, 40]. Most gut microbiota studies, including those on feeding paradigms use faecal samples as a representation of the gut, owing to convenient and non-invasive sampling procedures. However, as demonstrated in our study, the two body sites have distinct microbial communities and have different responses to changes in dietary composition and feeding patterns. Given the essential role of the microbial communities adhered to the colonic mucus layer in mediating host-microbiota interactions, examining the mucosal microbiota and their response to dietary interventions is crucial for understanding gut health.

### Fasting altered the colonic mucus layer *O*-glycosylation in a diet-dependent manner

To investigate the impact of fasting on gut mucin *O*-glycosylation, *O*-linked glycans were released from isolated mucin-2 glycoprotein (MUC2), the predominant mucin glycoprotein that makes the colonic mucus layer of mice. Released *O*-linked glycans were analysed using porous graphitised carbon liquid chromatography (PGC-LC) coupled with tandem mass spectrometry (MS/MS). A total of 43 unique mucin *O*-linked glycan structures, which were Core type 1 (Galβ1-3GalNAcαSer/Thr), 2 (GlcNAcβ1-6(Galβ1-3)GalNAcαSer/Thr) or 4 (GlcNAcβ1-6(GlcNAcβ1-3)GalNAcαSer/Thr), were identified (Table S3). These structures contained fucose (Fuc), sulfate, and N-acetylneuraminic acid (Neu5Ac) as terminal residues in varying combinations and degrees of substitution (Table S3). Mucin *O*-glycosylation is typically initiated with the attachment of N-acetylgalactosamine (GalNAc) residues to the hydroxyl group of Ser and Thr on the mucin protein backbone, which are then elongated into specific glycan core structures by the addition of galactose (Gal) and N-acetylglucosamine (GlcNAc) and can be terminated by terminal residues, including sulfate, Fuc and Neu5Ac [21]. Consistent with our observations, Core 1, 2, and 4 type *O-*linked glycans have been reported to be dominant in the colonic mucus layer of mice [15, 41].

There were significant impacts of the intermittent fasting regimen on the abundance of specific colonic mucin *O*-linked glycans (Fig. 4). Fucosylated glycan Hex1HexNAc1dHex1 (*m/z* 530.2), while abundant across all four groups, had significantly higher relative abundance in the HF-IF group compared to the HF group, with no significant difference between the NC and NC-IF groups. The relative abundances of five other glycans, Hex2HexNAc1dHex1 (*m/z* 692.1), isomers of Hex2HexNAc3NeuAc2 (*m/z* 766.8a and 766.8b), Hex2HexNAc3dHex2 (*m/z* 1244.5c) and Hex2HexNAc3NeuAc1dHex2 (*m/z* 1389.5a), were lower in the HF-IF compared to the HF group. This demonstrates a clear impact of fasting on colonic mucin *O*-glycan composition, particularly on Core 1 and Core 4 type glycan structures of mice fed the high-fat diet. A different set of glycans, which were Core 2 type structures, were found to be significantly different between the NC and NC-IF groups. In particular, the fucosylated glycan Hex1HexNAc2dHex1 (*m/z* 733.3) was observed at a higher relative abundance while sulphated glycan Hex1HexNAc2dHex1S (*m/z* 813.2) and difucosylated Hex2HexNAc2dHex2 (*m/z* 1041.4) had a lower relative abundance in the NC-IF group compared to the NC group. Our results demonstrate a clear and diet-specific impact of intermittent fasting on colonic mucin *O*-glycosylation; fasting had a greater impact on Core 1 and 4 type *O*-linked glycan structures in the colonic mucosa of mice fed the high-fat diet, while it had a higher impact on Core 2 type *O*-glycan structures when mice were fed the normal chow. Furthermore, changes in the HF-IF group also suggested an amelioration of some high-fat diet-induced changes in mucin *O*-glycosylation. The relative abundance of fucosylated and sialylated glycans Hex2HexNAc1dHex1 (*m/z* 692.1), Hex2HexNAc3NeuAc2 (*m/z* 766.8b) and Hex2HexNAc2dHex1 (*m/z* 895.3) were significantly higher in the HF group while the difucosylated glycan Hex2HexNAc2dHex2 (*m/z* 1041.4) was observed at a lower relative abundance compared to the NC. While fasting did not reverse all high-fat diet-induced changes in *O*-glycosylation, a fucosylated (*m/z* 692.1) and a doubly sialylated (*m/z* 766.8b) glycan, which were observed at a higher relative abundance in the HF group, had a lower relative abundance in the HF-IF group.

**Figure 4.**
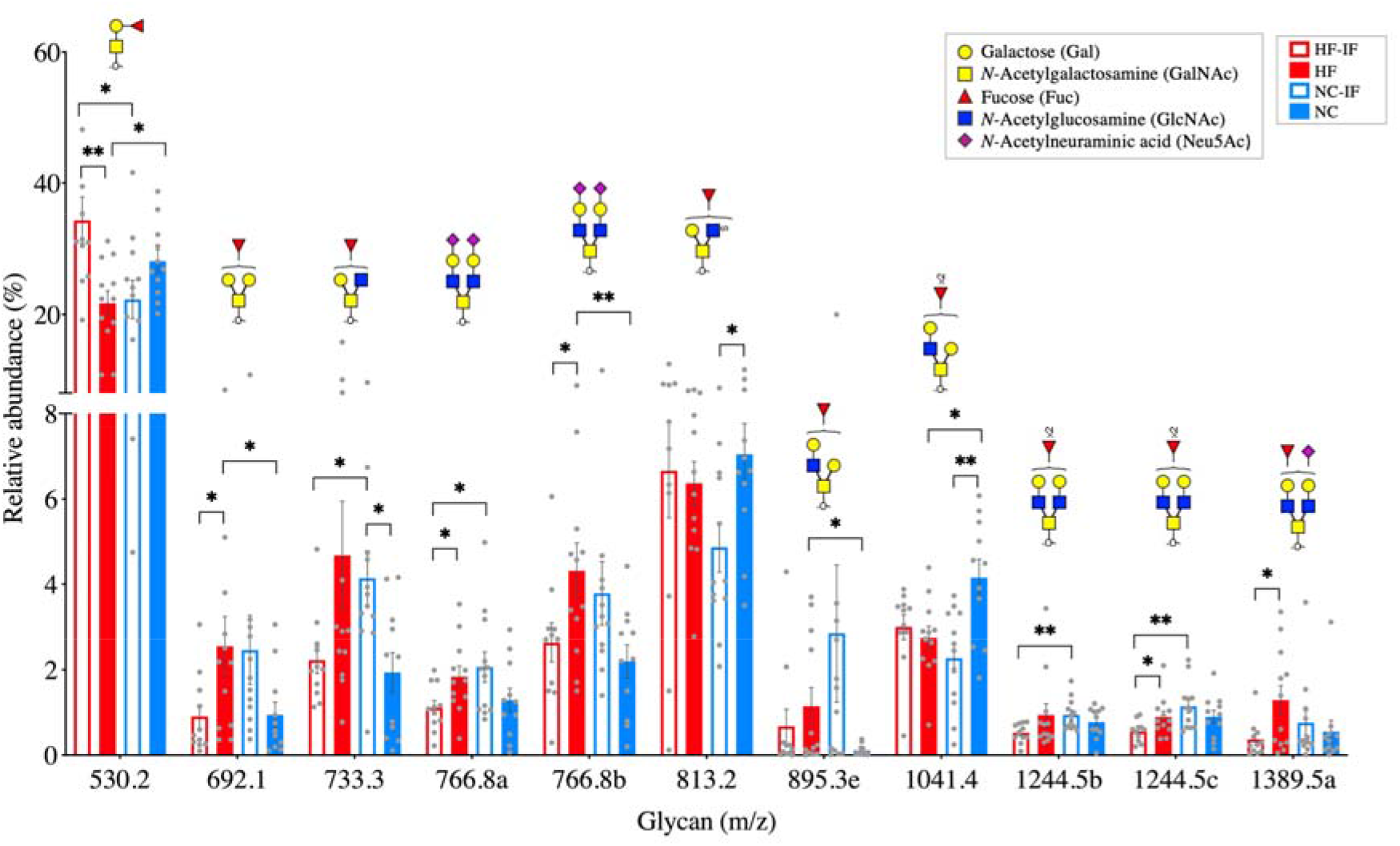
Impact of intermittent fasting on the relative abundance of colonic mucin *O*-linked glycan structures. *O*-glycans were released from the colonic mucosa of mice in each of the four treatment groups: high-fat diet with intermittent fasting (HF-IF) or *ad libitum* (HF) and normal chow with intermittent fasting (NC-IF) or *ad libitum* (NC). Glycans are listed by their negative ion mass to charge (m/z) with structural isomers distinguished alphabetically. Data are shown as mean ± SD for each dietary group. Statistical significance was determined using Student’s t-test between groups (** *P* < 0.01 and * *P* < 0.05). A complete list of glycans identified and their abundances are provided in Table S3.

The colonic mucus layer is crucial not only in providing physical and chemical separation between the host and colonic lumen, but also in maintaining host-microbiota interactions. As the predominant component of the colonic mucosa, mucin proteins and their *O*-linked glycans are vital in shaping the microbial populations in the gut [23]. Alterations in mucin *O*-linked glycans, particularly terminal residues, such as N-acetylneuraminic acid, fucose and sulfate have profound impacts on gut health, including a role in maintaining mucus layer structural integrity [42, 43]. For example, N-acetylneuraminic acid has been shown to contribute to regulating rheological properties of the mucus layer promoting rigidity and viscosity [43]. Despite the critical role of *O*-linked glycans in regulating the gut environment, the impact of dietary changes on *O*-glycosylation remains relatively unknown. To our knowledge, our study is the first to use modern glycomic techniques to demonstrate a significant impact of intermittent fasting on colonic mucin *O*-glycosylation profile. Future studies designed to examine other markers of gut health, including mucus layer thickness, rigidity, and viscosity will be useful extensions in gaining further insights into the impact of fasting on the gut and its mucus layer.

### Alterations in mucin *O*-linked glycans associated with changes in specific mucosal bacteria

Pairwise correlation analysis between the relative abundance of bacterial ASVs and the relative abundance of mucin *O*-linked glycans identified how changes in the mucosal microbiota associate with changes in colonic mucosal *O*-glycosylation upon fasting. Associations between mucosal bacteria and *O*-linked glycans that were significantly different in the HF-IF and NC-IF groups compared to the HF and NC groups, respectively, were used to construct separate correlation networks for each diet (Fig. 5). The higher relative abundance of glycan *m/z* 530.2 in the HF-IF group showed a negative correlation with ASVs in the bacterial family *Lachnospiraceae*, while these ASVs showed positive correlations with the glycan *m/z* 692.1 (Fig. 5A). This may suggest a preference in members of *Lachnospiraceae* for glycan *m/z* 692.1 over *m/z* 530.2. Both glycans are Core 1 type structures. Glycan *m/z* 692.1 contains an additional terminal hexose (Gal) residue attached to its GalNac (Fig. 5A). Two ASVs in the family *Muribaculaceae*, which showed a negative correlation with glycan *m/z* 530.2, had a positive correlation with two isomers of glycan *m/z* 766.8. These isomers are Core 4 type structures that are elongated with two sialic acid terminal residues. Upon examination of microbe-glycan associations in the NC-IF group, we observed positive correlations of ASVs in *Christensenellaceae* and *Mollicutes* RF39 with glycan *m/z* 813.2, while they had negative correlations with glycan *m/z* 733.3. Glycan *m/z* 733.3 is the precursor of glycan *m/z* 813.2 synthesis, which has an additional sulfate residue (Fig. 5B). Our results suggest that fasting-mediated changes in colonic mucin *O*-glycosylation could have a profound impact on the mucosal microbiota and *vice versa*, potentially through their bidirectional interactions that are essential for regulating gut health and host-microbiota interactions.

**Figure 5.**
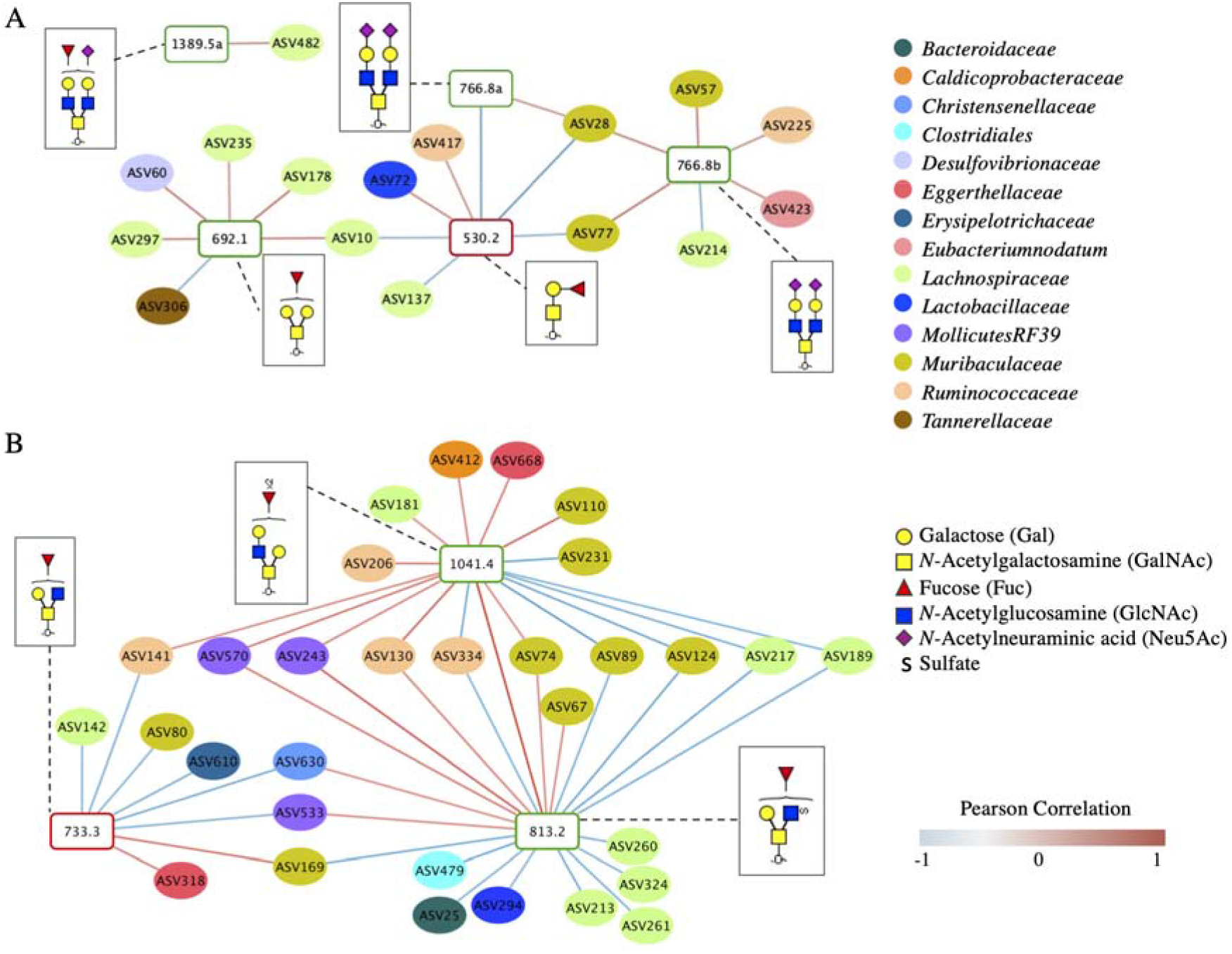
Networks showing correlations between amplicon sequence variants (ASVs) in the colonic mucosal microbiota and mucin *O*-glycans. The associations between the relative abundance of ASVs and relative abundance of *O*-glycans were identified using pairwise correlation analysis, significant correlations (*P* < 0.05) were used to construct networks. Only the ASVs and glycans with significantly different abundances in the **(A)** high-fat diet with intermittent fasting (HF-IF) group compared to the high-fat diet *ad libitum* (HF) group and **(B)** normal chow with intermittent fasting (NC-IF) group compared to the normal chow *ad libitum* (NC) group were included in the analysis. The ASVs are shown in colour-coded dots based on the bacterial family. Glycans are listed by their negative ion mass to charge (m/z) with structural isomers distinguished alphabetically. The increase or decrease in the abundance of glycans are indicated with a red or green outline, respectively. A positive or negative Pearson correlation coefficient is presented by a red and blue line, respectively, the intensity of the colour denotes the strength of the correlation.

Mucin *O*-glycans provide binding targets for gut bacterial adhesion [44], serve as an alternate energy source for gut microbiota [45, 46] and regulate gut microbial communities through the application of selective pressures [47, 48], all of which can shape the microbiota composition and functions [15, 46, 49]. Specific gut bacteria contain cell surface adhesion sites and lectins (glycan receptor proteins), which are essential for binding to colonic mucin *O*-linked glycans, while other members can digest *O*-glycans [50, 51]. Gut bacteria use a range of enzymes, including sulfatases and glycosyl hydrolases (GHs) for metabolising mucin *O*-linked glycans and using them as an energy source [52]. On the other hand, gut microbial activities, particularly changes in the production of microbial metabolites, such as acetate, have been shown to modulate *O*-glycosylation by affecting the expression of host glycosyl transferases, which are enzymes involved in the synthesis of mucin *O*-linked glycans [53, 54]. Correlations observed in our study could potentially reflect these microbiota-glycan interactions. Bacterial ASVs in the families *Lachnospiraceae, Muribaculaceae*, and *Rumminococceae* that showed strong correlations with mucin *O*-linked glycans are known to express proteins and enzymes that could potentially recognise, bind, and digest these glycans. For example, members in *Lachnospiraceae* encode for carbohydrate binding proteins and GH family enzymes, such as galactosidases (GH98) [35, 55]. *Muribaculaceae* is a member of the phylum *Bacteroidetes* in which specific members have been shown to express a range of GH family enzymes, including sialidases (GH33) [56]. This is consistent with our observations where ASVs in *Lachnospiraceae* and *Muribaculaceae* showed positive correlations with glycans containing galactose (Gal) and N-Acetylneuraminic acid (Neu5Ac, sialic acid) terminal residues, respectively. While the potential of these bacteria to bind and/or digest glycans could explain the correlations observed in our study, direct evidence is currently unavailable due to a lack of culture isolates and individual genome analysis of these specific ASVs. Genomic and biochemical investigations on the individual bacteria and how they interact with mucin *O*-glycans are essential to confirm the biological mechanisms of these associations.

The intimate interactions between mucin *O*-linked glycans in the colonic mucus layer and mucosal microbiota are not only important in maintaining gut health but also in shaping the gut microbiota and its associations with the host. While it is evident that intermittent fasting promotes gut and host health, there are no reports to date on how fasting affects the colonic mucus layer composition and mucosal microbiota. Our study provides the groundwork for a comprehensive understanding on how intermittent fasting influences colonic mucin *O*-glycans and their associations with specific mucosal bacteria. Altering or supplementing mucin *O*-glycan compositions in the colon could be a targeted approach for gut microbiota modulation [57]. The outcomes of our study suggest that intermittent fasting could be used to achieve this in a diet-specific manner.

## Conclusions

Consuming a high-fat diet with intermittent fasting was associated with a significantly lower body weight, improved glucose tolerance and reduced plasma resistin and leptin levels compared to the groups with *ad libitum* access to the high-fat diet. These improvements were observed despite the similar caloric intake between these two groups. Fasting had no significant impact on the physiology of mice fed a normal chow. Conversely, intermittent fasting had clear and distinct impacts on the mucosal and faecal microbial community structure and composition and colonic mucin *O*-glycosylation of mice fed either the high-fat diet or normal chow. However, the precise nature of changes was specific to each diet, demonstrating a strong link of dietary composition with the beneficial effects of short-term exposure to intermittent fasting. Furthermore, diet-specific associations were identified between specific colonic mucosal bacteria and mucin *O*-glycans, which suggested an impact of fasting on how the gut microbiota interact with the colonic mucus layer.

This is the first comprehensive investigation providing insight into the influence of intermittent fasting on the mucosal microbiota and colonic mucin *O*-glycosylation. The intimate associations between the gut microbiota and mucin glycans are critical but remain largely unexplored. Our study provides significant insight into how changes to the dietary composition and feeding patterns affect the interactions between specific mucosal bacteria and mucin *O*-glycans.

## Materials and Methods

### Animal trial and sample collection

All protocols and procedures were reviewed and approved by the Animal ethics committee, Macquarie University, Australia (ARA 2018/003).

A total of 47, six- to seven-week-old male C57BL/6J mice were obtained from the Animal Resource Centre, WA, Australia. Three animals per cage were cohoused under standard and monitored conditions: temperature (20-26 °C), humidity (40-60%), light and dark cycle (12 hour-12 hour) and with *ad libitum* access to water throughout the experiment. Following one week acclimatisation on a normal chow (14.0 kJg^-1^ digestible energy, 12% of total digestible energy from lipids), a group of mice were randomised into a high-fat diet (19.0 kJg^-1^ digestible energy, 43% of total digestible energy from lipids). Each dietary group was maintained with either *ad libitum* or with intermittent fasting, resulting in four treatment groups: normal chow *ad libitum* (n=11, NC), normal chow with intermittent fasting (n=12, NC-IF), high-fat diet *ad libitum* (n=12, HF) and high-fat diet with intermittent fasting (n=12, HF-IF). The two groups subjected to intermittent fasting did not have access to feed for 24-hour periods twice a week on two non-consecutive days (Monday and Thursday) for 12 weeks, these animals had *ad libitum* access to food on the remaining five days each week. Details of the experimental design are provided in Figure S1. All experimental diets were produced by Speciality feeds, WA, Australia. Nutritional information and ingredients are provided in Table S1.

Individual body weight was measured at least three time a week: Monday (72 hours after fasting), Wednesday (24 hours after fasting) and Friday (immediately after fasting). Food intake per cage was measured weekly. Faecal samples were aseptically collected from individual mice at week 0 and 12 and stored at -80°C prior to subsequent microbiota analyses.

Blood (100 µL) was collected from the tail vein at week 0 and 12 after overnight fasting with *ad libitum* access to water. These samples were gently mixed with EDTA (final concentration 4 mM and pH 7.0). The plasma was separated by centrifugation at 1000 x g for 10 min at 4 °C and stored□at -80 °C prior to further analyses.

Mice were euthanised at week 12 by cervical dislocation. Body organs were excised, the liver, caecum and epididymal fat pads were weighed. A section of the liver was fixed in 4% (v/v) formaldehyde and stored at 4 °C prior to histological analysis. The colonic content was removed, and the colon was cleaned with 1x phosphate buffered saline (PBS) before storing at -80 °C.

### Intraperitoneal glucose tolerance tests

Mice were fasted for six hours during the light cycle prior to performing intraperitoneal glucose tolerance tests (IPGTTs) in week 4 and 11. Initial blood glucose levels were measured from the tail vein using a Freestyle Lite blood glucose monitoring system (Abbott Pty Ltd, Australia). Following injection of glucose (2.0 gkg^-1^, intraperitoneally), blood glucose levels were measured at 15, 30, 60, 90 and 120 minutes.

### Histological analysis

Liver tissues fixed in in 4% formaldehyde were transferred into a 15% (w/v) sucrose solution in 1x PBS and incubated at 4 °C for 6 hours or until saturated. The tissues were transferred to a 30% (w/v) sucrose solution in 1x PBS and incubated at 4 °C overnight or until saturated. Tissues were dehydrated, embedded in the optimal cutting temperature (OCT) compound, and sliced (6 µm) using a Cryostat CM3050 S (Leica, Australia) according to the manufacturer’s instructions. The slices were stained with hematoxylin and eosin (H and E) for histological examination using an Olympus BX 63 microscope.

### Quantification of circulating plasma inflammatory, obesity and diabetes markers

The blood plasma cytokine, and obesity and diabetes markers were quantified using Bio-Plex Pro™ mouse cytokine (M60000007A), and obesity and diabetes (171F7001M) assay kits, respectively, according to the manufacturer’s instructions (Bio-Rad, Australia).

### 16S rRNA gene amplicon sequencing and bioinformatics analysis

The colonic mucus was obtained by vacuum suction from the entire length of the colon, for this, the whole colon was opened longitudinally, gently flushed with 100 µL of 1x PBS and total mucus was extracted using a vacuum. Total community DNA was isolated from colonic mucus samples (n=47) and faecal samples collected at week 0 and 12 (n= 94) using a FastDNA spin kit (MP Biomedicals, Australia) according to the manufacturer’s instructions. The V4 region of the 16S rRNA gene was amplified using Five prime hot master mix (VWR, Australia) using primers with custom barcodes, 515F (5□-GTGCCAGCMGCCGCGGTAA-3□) and 806R (5□-GGACTACHVGGGTWTCTAAT-3□). The resulting amplicons were quantified (Quant-iT™ PicoGreen^®^ Invitrogen, Australia), equal molar amounts of barcoded amplicons from each sample were pooled, gel purified using a Wizard^®^ SV gel and PCR clean up system (Promega, Australia) and sequenced using an Illumina MiSeq platform at the Ramaciotti Centre for Genomics, Australia.

Raw sequence data was processed using Quantitative Insights Into Microbial Ecology software (QIIME2 version 2018.6) [58]. Demultiplexed paired-end reads were quality filtered using the q2□demux plugin (median q<30), followed by denoising with DADA2 [59]. Amplicon sequence variants (ASVs) were aligned with mafft [60] (via q2□alignment) and used to construct a phylogeny with fasttree2 [61] (via q2□phylogeny). The generated ASVs were assigned to taxonomy at 99% similarity using the q2□feature□classifier [62] classify□sklearn nai□ve Bayes taxonomy classifier against the SILVA 132 database [63] of the 515F/R806 region of the 16S rRNA gene. A total of 7, 925,219 reads (mean 56,207 ± 20,781) were obtained after quality filtering, each of the 141 samples were rarefied at 20,000 reads prior to statistical analyses. Two random samples that failed to meet this requirement were eliminated from further analyses.

Statistical analysis of the gut microbiota sequencing data was conducted using PRIMER-7 software package [64]. Non-metric multidimensional scaling (nMDS) plots were constructed based on Bray-Curtis similarity matrices of Log (x + 1) transformed abundance of the ASVs. Permutational Multivariate Analysis of Variance (PERMANOVA) tests with 9999 permutations were conducted to investigate differences in the microbial community structure. The Shannon diversity and Simpson’s evenness indices for each sample was determined using the PRIMER-7 software package.

Distinct ASVs between treatment groups were identified using the Linear Discriminant Analysis (LDA) effect size (LEfSe) method (online Galaxy Version 1.0) [65] using default parameters. Treatment groups were used as classes of subjects with no subclasses. LEfSe analyses were performed using default parameters: Kruskal-Wallis test among classes (*P* < 0.05), Wilcoxon test between classes (*P* < 0.05) and the threshold on the logarithmetic LDA score for discriminative features > 2.0.

### Mucin extraction and *O*-glycan characterisation

Mucins were obtained from the extracted colonic mucus by precipitation with 6M guanidine hydrochloride. *O-*glycans were released from the mucins by reductive β-elimination and subjected to PGC-LC-ESI-MS/MS analysis according to a previously established protocol [66] using a Dionex Ultimate 3000 RS HPLC system coupled to a Thermo LTQ Velos Pro (Thermo Scientific, USA). *O-*glycan compositions were calculated from the masses using GlycoMod (https://web.expasy.org/glycomod/) and glycan structures were assigned by manual interpretation of the tandem MS fragmentation spectra. Glycan peaks were quantified by relative abundance using Skyline (Version 3.7.0.11317, MacCoss Lab, UW) for assisted peak picking, generation of extracted ion chromatograms (EICs) and integration of EIC peak areas. Glycan structures were drawn using GlycoWorkbench 2 (Version 2.1) using SNFG nomenclature.

### Statistical analysis

D’Agostino-Pearson normality tests were performed prior to determining the statistical significance of differences observed in the body weight, feed intake, IPGTT, concentration of biomarkers, organ weight and microbiota alpha diversity. Either Kruskal-Wallis test with Dunn’s multiple comparisons test or Tukey’s multiple comparisons test was performed between treatment groups to determine the statistical significance of differences using GraphPad Prism (version 8.4) software (GraphPad Software, La Jolla California, USA). The significance of the differences observed in the abundance of mucin *O*-glycans between treatment groups was determined using Student t-tests.

### Correlation network analysis

The associations between the relative abundance of mucus adhered bacterial ASVs and colonic mucin *O-*linked glycans were identified using pairwise correlation analyses using Hmisc R package [67]. Only bacterial ASVs and glycans with significantly different abundances in the HF-IF and NC-IF groups compared to the HF and NC groups, respectively, were used for the analysis. Statistically significant correlations (*P* < 0.05) between the ASVs and *O*-glycan structures were used to generate correlation networks using Cytoscape software (Version 3.6.1).

## Data availability

The 16S rRNA gene sequence data generated and analysed during the current study are available in the GenBank Sequence Read Archive database under accession number PRJNA788645 https://dataview.ncbi.nlm.nih.gov/object/PRJNA788645?reviewer=nlogfk4edjruupdjvaid2fcrg6

## Authors’ contributions

HKAHG, NHP and ITP designed the research plan. HKAHG performed the animal trial; analysed host physiological parameters and the microbiota; and conducted bioinformatics, statistical and correlation network analyses. AMMS performed glycomics. AMMS, KN and RWWC analysed glycomics data. HKAHG wrote the manuscript, which was edited by RWWC, KN, AMMS, NHP and ITP. All authors read and approved the final manuscript.

## Acknowledgements

This work was supported by the National Health and Medical Research Council (GNT1127292) and Australian Research Council Centres of Excellence (CE200100029 and CE140100003) funded by the Australian government. KN was supported by a joint Macquarie University/Thermo Scientific scientist sponsorship.

We thank Carlie Crawford at the Central Animal House Facility, Macquarie University for providing technical support.

## Supplementary Figures and Tables

**Figure S1.**
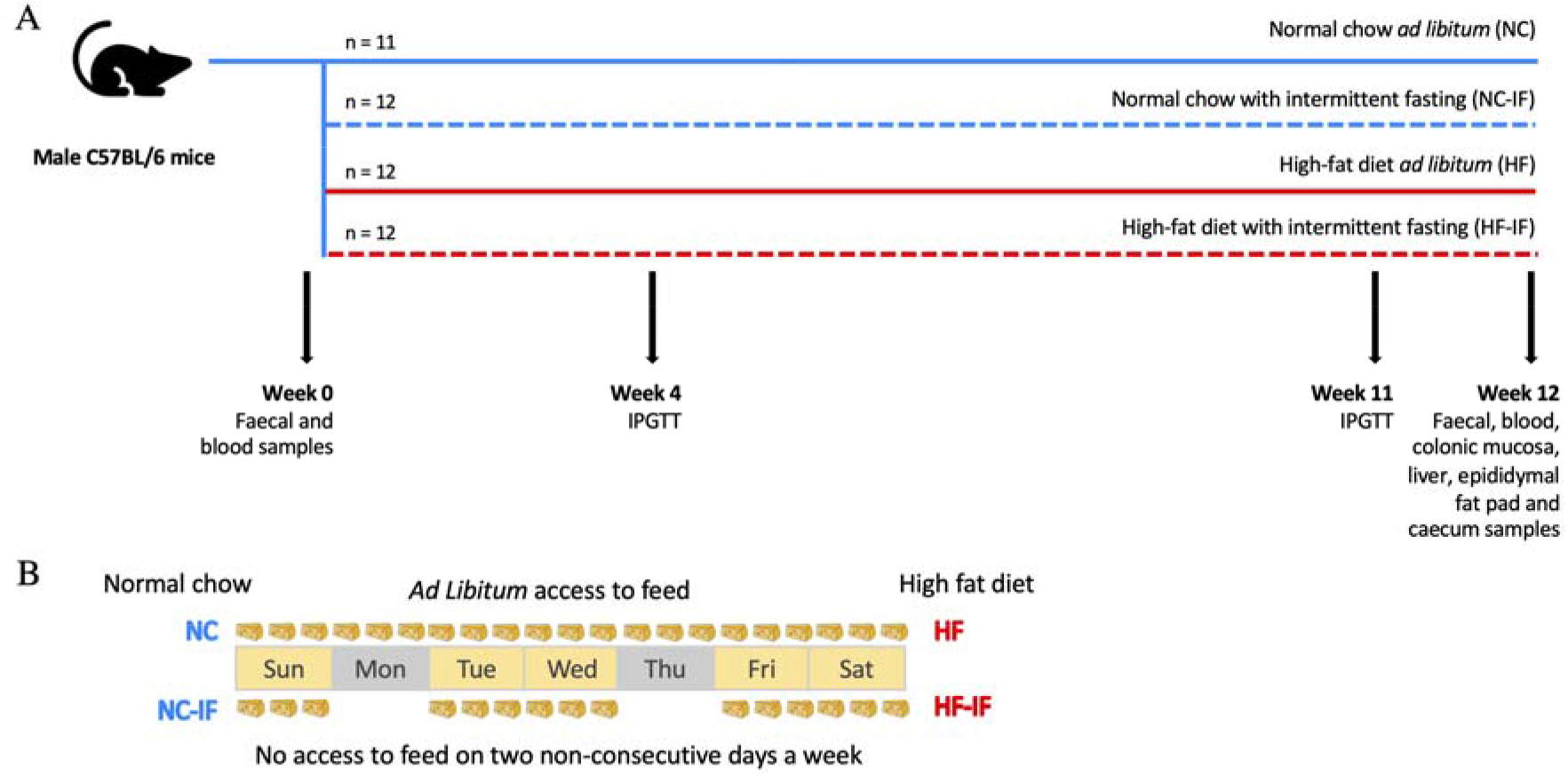
**(A)** Experiment timeline. A group of 47 male 6-7 weeks old C57BL/6J mice were randomised into one of four groups: high-fat diet with intermittent fasting (HF-IF) or *ad libitum* (HF), or normal chow with intermittent fasting (NC-IF) or *ad libitum* (NC). Individual body weight and feed intake per cage were measured thrice and once a week, respectively. Intraperitoneal glucose tolerance tests (IPGTTs) were conducted at weeks 4 and 11. Faecal and blood samples were collected at weeks 0 and 12. The colonic mucosa, liver, epidydimal fat pads and caecum were harvested upon euthanasia. **(B)** Schedule of the intermittent fasting regimen. The HF-IF and NC-IF groups had no access to feed for 24 hours each on two non-consecutive days a week (Monday and Thursday), this was continued for 12 weeks. Feed was provided *ad libitum* on non-fasting days. HF and NC groups had *ad libitum* access to their respective feed. All groups had *ad libitum* access to water throughout the experiment.

**Figure S2.**
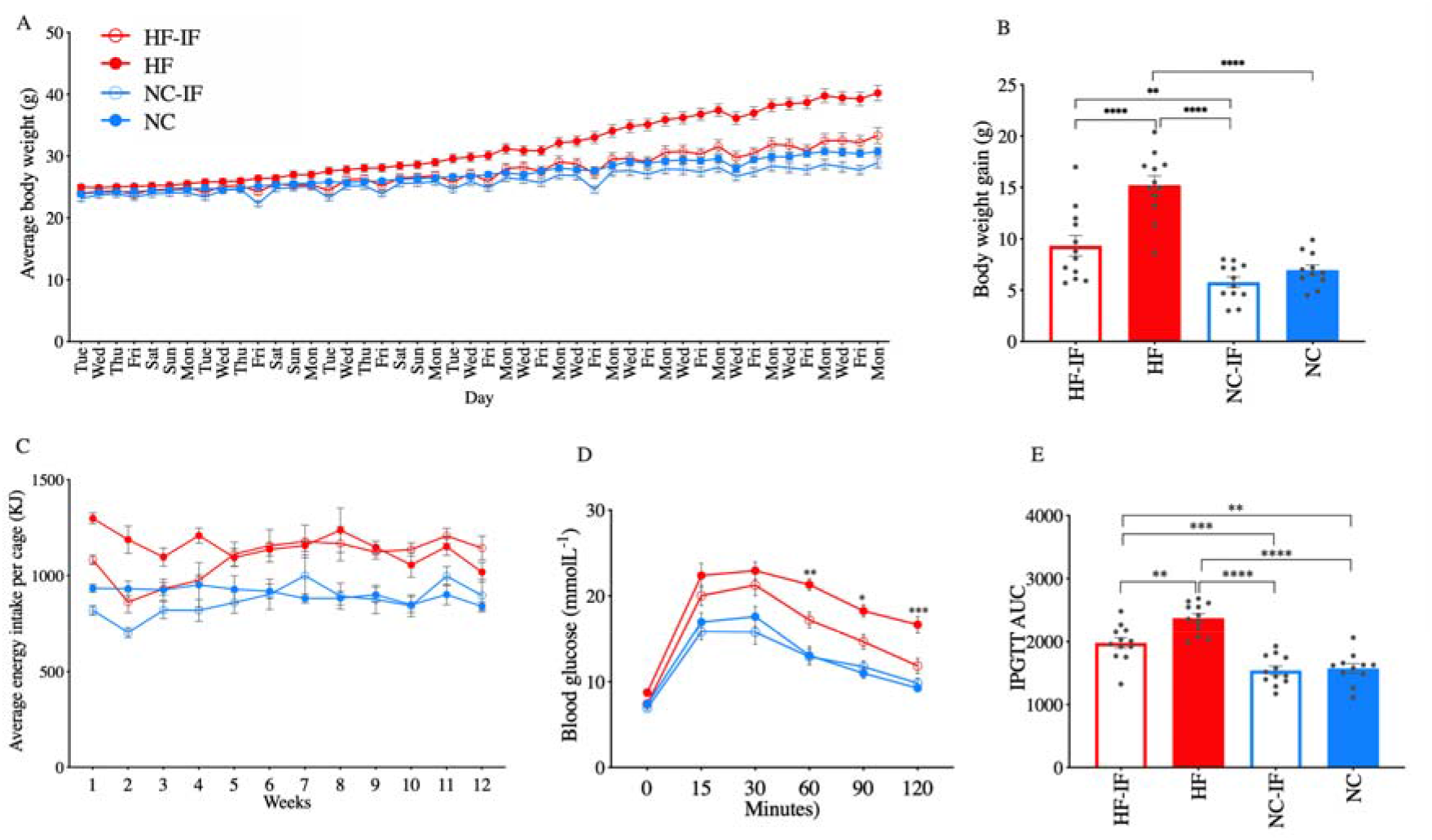
Impact of intermittent fasting on body weight and glucose tolerance. **(A)** The body weight per mouse was measured at least on three days a week for 12 weeks; **(B)** body weight gain from week 0 to week 12 per mouse was determined; **(C)** weekly energy intake per cage was calculated; **(D)** blood glucose levels and **(E)** the area under the curve (AUC) of the intraperitoneal glucose tolerance test (IPGTT) conducted at week 11 are shown. Data are presented as mean ± SD for each treatment group: high-fat diet with intermittent fasting (HF-IF) or *ad libitum* (HF) and normal chow with intermittent fasting (NC-IF) or *ad libitum* (NC). Significance was determined based on two-way ANOVA with Tukey’s multiple comparison tests or Kruskal-Wallis test with Dunn’s multiple comparison tests where appropriate (**** *P* < 0.0001, *** *P* < 0.001, ** *P* < 0.01 and * *P* < 0.05).

**Figure S3.**
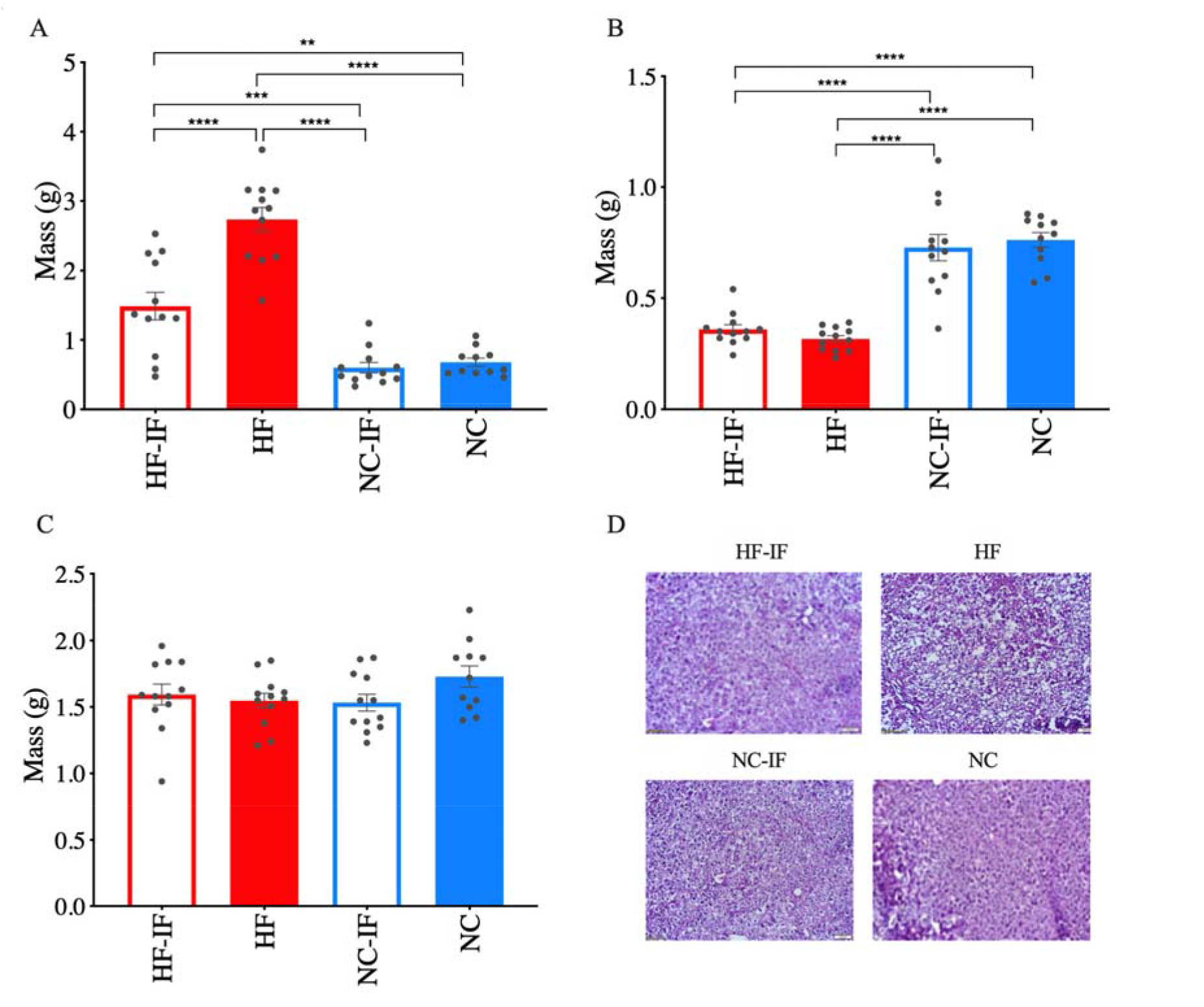
Mass of specific body organs and hematoxylin and eosin (H and E) staining of liver sections harvested from mice at week 12. **(A)** Epididymal white adipose tissue, **(B)** caecum, and **(C)** liver masses are shown. Mean ± SD values are shown for each treatment group: high-fat diet with intermittent fasting (HF-IF) or *ad libitum* (HF) and normal chow with intermittent fasting (NC-IF) or *ad libitum* (NC). Significance was determined using Kruskal-Wallis test with Dunn’s multiple comparison tests (**** *P* < 0.0001, *** *P* < 0.001, ** *P* < 0.01 and * *P* < 0.05). **(D)** Representative microscopic images (x10) of the liver tissues with H and E staining. Liver samples excised from mice at week 12 were fixed in 4% (v/v) formaldehyde prior to processing for microscopic examination.

**Figure S4.**
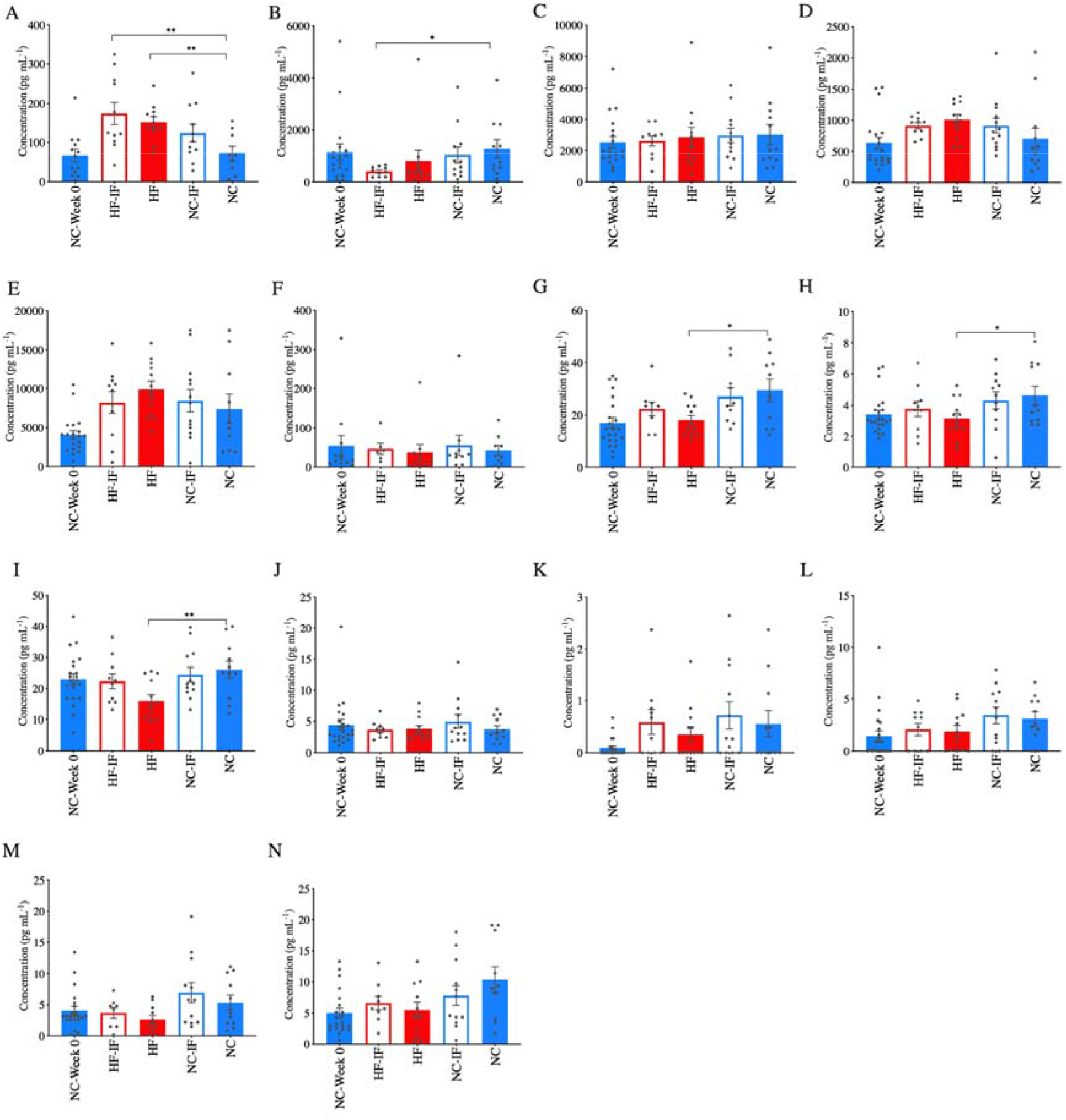
Concentrations of specific cytokines and biomarkers of diabetes and obesity in blood plasma collected at weeks 0 and 12. Mean ± SD values are shown for **(A)** gastric inhibitory polypeptide (GIP), **(B)** glucagon, **(C)** insulin, **(D)** plasminogen activator inhibitor-1 (PAI1), **(E)** ghrelin, **(F)** glucagon-like peptide-1 (GLP-1), **(G)** tumour necrosis factor (TNF) alpha, **(H)** interleukin (IL)-2, **(I)** IL-10, **(J)** IL-5, **(K)** IL-4, **(L)** Granulocyte-macrophage colony-stimulating factor (GM-CSF), **(M)** IL-1B and **(N)** Interferon (IFN) gamma. Kruskal-Wallis test with Dunn’s multiple comparison tests was used to determine the significance of differences (** *P* < 0.01 and * *P* < 0.05) observed between treatment groups: high-fat diet with intermittent fasting (HF-IF) or *ad libitum* (HF) and normal chow with intermittent fasting (NC-IF) or *ad libitum* (NC).

**Figure S5.**
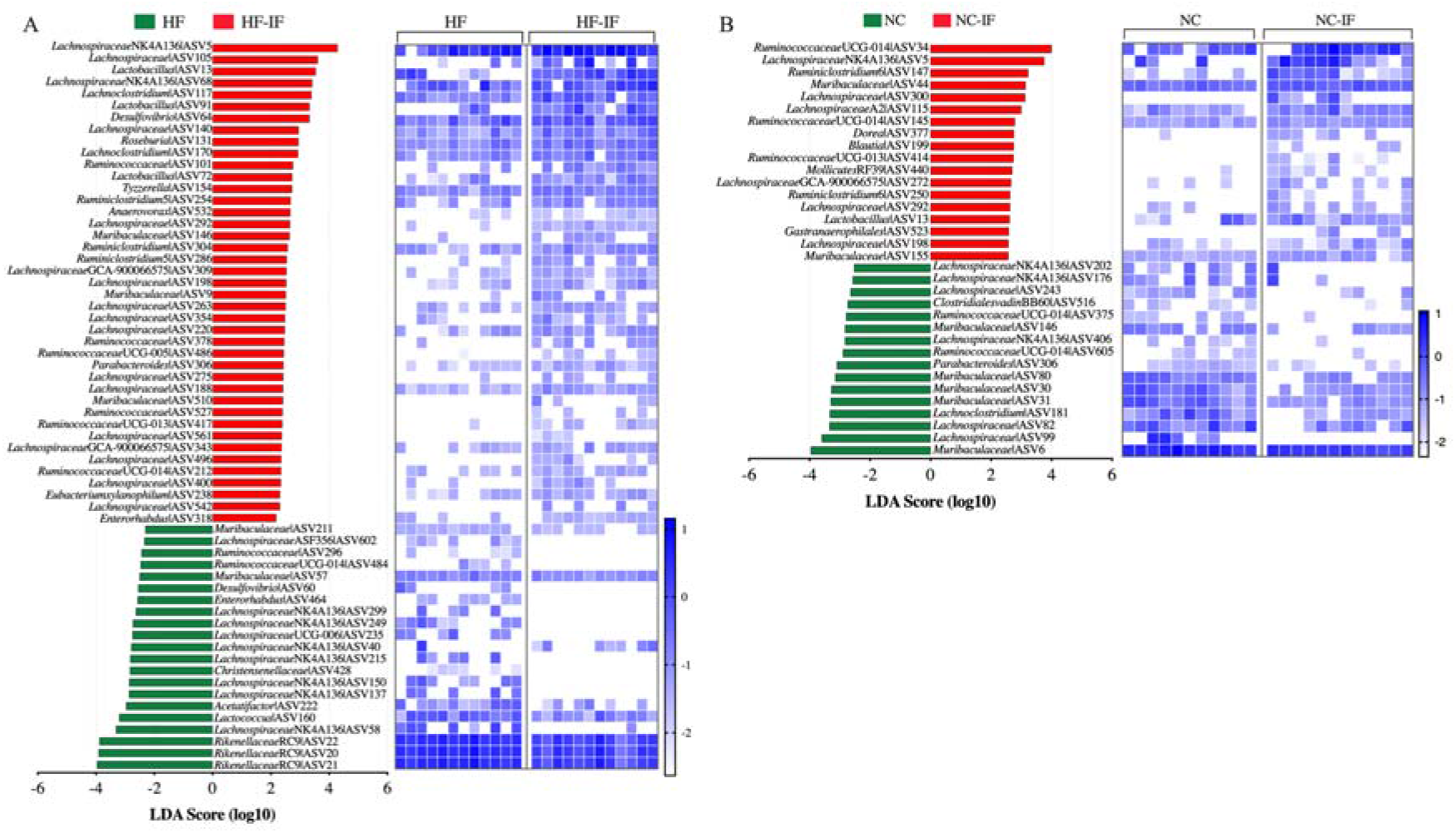
Effect of intermittent fasting on the abundance of amplicon sequence variants (ASVs) in the faecal microbiota. Data were obtained using linear discriminant analysis effect size (LEfSe) method through comparing the abundance of ASVs between treatment groups at week 12. **(A)** High-fat diet *ad libitum* vs intermittent fasting (HF vs HF-IF) and **(B)** normal chow *ad libitum* vs intermittent fasting (NC vs NC-IF). The histograms show linear discriminant analysis (LDA) scores computed for each ASV with significantly different abundance between groups. The unpaired heatmaps show the relative abundance (Log_10_ transformed) of the ASVs, rows correspond to the abundance of the ASVs, and columns correspond to individual mice in each dietary group. Blue and white denote the highest and lowest relative abundance, respectively, as shown in the legend.

**Figure S6.**
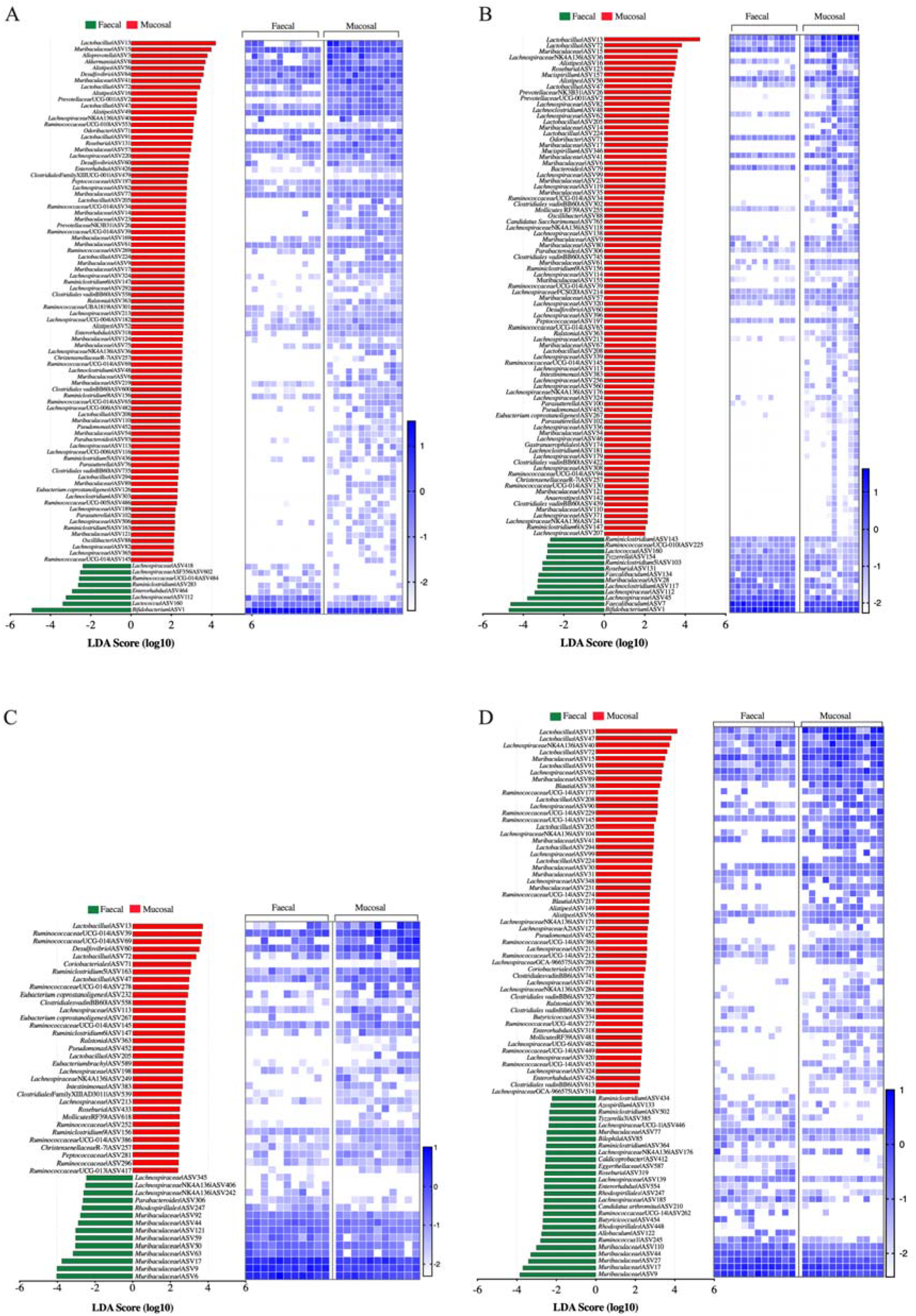
Each diet influenced the abundance of amplicon sequence variants (ASVs) in the faecal and mucosal microbiota differently. Data were obtained using linear discriminant analysis effect size (LEfSe) method through comparing the mucosal and faecal microbiota of mice in the **(A)** high-fat diet *ad libitum* (HF), **(B)** high-fat diet with intermittent fasting (HFIF), **(C)** normal chow *ad libitum* (NC) and **(D)** normal chow with intermittent fasting (NC-IF) groups. The histograms show linear discriminant analysis (LDA) scores computed for each ASV with significantly different abundance between groups. The unpaired heatmaps show the relative abundance (Log_10_ transformed) of the ASVs, rows correspond to the abundance of the ASVs, and columns correspond to individual mice in each dietary group. Blue and white denote the highest and lowest relative abundance, respectively, as shown in the legend.

**Table S1** Nutritional information and ingredients of the high-fat diet and normal chow used in the study.

**Table S2** Amplicon sequence variants (ASVs) that were found to have significantly different abundances between the high-fat diet *ad libitum* (HF) and normal chow *ad libitum* (NC) groups.*

*Data was obtained using linear discriminant analysis effect size (LEfSe) through comparing the abundance of ASVs in the **(A)** mucosal microbiota of the high-fat diet *ad libitum* (HF) vs normal chow *ad libitum* (NC) groups, and (B) faecal microbiota of the HF vs NC groups. Taxonomic assignment of each ASVs is shown with the linear discriminant analysis (LDA) score.

**Table S3** The relative abundance of *O-*linked glycans of the mucin-2 glycoprotein isolated from the mouse colon.*

*HF- high-fat diet *ad libitum*, HF-IF- high-fat diet with intermittent fasting, NC- normal chow *ad libitum*, NC-IF- normal chow with intermittent fasting, Values are provided as % relative abundance for each mouse. Glycans are listed by their negative ion mass to charge (m/z) with structural isomers distinguished alphabetically.

